# Multiplexed competition in a synthetic squid light organ microbiome using barcode-tagged gene deletions

**DOI:** 10.1101/2020.08.24.265777

**Authors:** Hector L. Burgos, Emanuel F. Burgos, Andrew J. Steinberger, Garret Suen, Mark J. Mandel

**Affiliations:** Department of Medical Microbiology and Immunology, University of Wisconsin-Madison, Madison, WI USA; Department of Bacteriology, University of Wisconsin-Madison, Madison, WI USA

**Keywords:** Barcode sequencing, amplicon sequencing, sequence-tagged gene deletions, *Vibrio fischeri*, *Aliivibrio fischeri*

## Abstract

Beneficial symbioses between microbes and their eukaryotic hosts are ubiquitous and have widespread impacts on host health and development. The binary symbiosis between the bioluminescent bacterium *Vibrio fischeri* and its squid host *Euprymna scolopes* serves as a model system to study molecular mechanisms at the microbe-animal interface. To identify colonization factors in this system, our lab previously conducted a global transposon insertion sequencing (INSeq) screen and identified over 300 putative novel squid colonization factors in *V. fischeri*. To pursue mechanistic studies on these candidate genes, we present an approach to quickly generate barcode-tagged gene deletions and perform high-throughput squid competition experiments with detection of the proportion of each strain in the mixture by barcode sequencing (BarSeq). Our deletion approach improves on previous techniques based on splicing-by-overlap extension PCR (SOE-PCR) and *tfoX*-based natural transformation by incorporating a randomized barcode that results in unique DNA sequences within each deletion scar. Amplicon sequencing of the pool of barcoded strains before and after colonization faithfully reports on known colonization factors and provides increased sensitivity over colony counting methods. BarSeq enables rapid and sensitive characterization of the molecular factors involved in establishing the *Vibrio*-squid symbiosis and provides a valuable tool to interrogate the molecular dialogue at microbe-animal host interfaces.

**Importance:** Beneficial microbes play essential roles in the health and development of their hosts. However, the complexity of animal microbiomes and general genetic intractability of their symbionts have made it difficult to study the coevolved mechanisms for establishing and maintaining specificity at the microbe-animal host interface. Model symbioses are therefore invaluable for studying the mechanisms of beneficial microbe-host interactions. Here we present a combined barcode-tagged deletion and BarSeq approach to interrogate the molecular dialogue that ensures specific and reproducible colonization of the Hawaiian bobtail squid by *Vibrio fischeri*. The ability to precisely manipulate the bacterial genome, combined with multiplex colonization assays, will accelerate the use of this valuable model system for mechanistic studies of how environmental microbes—both beneficial and pathogenic—colonize specific animal hosts.

## Introduction

Beneficial symbioses are ubiquitous in the environment and have substantial impacts on the health and development of animal hosts. In animals, symbionts can affect host organ morphogenesis, immune system development, reproduction, susceptibility to disease, and even behavior (1–4). In humans, the gut, skin, lungs, and urogenital tract all have specific microbiomes for which their dysbiosis has been associated with disease (5–8). It is clear that molecular communication between animal hosts and their microbial partners leads to selection and retention of the cognate microbiome: while many microbes are obtained from the environment, the composition of mature microbiomes is often largely stable and resilient within members of a host species (9, 10). While microbial communities have been characterized using metagenomic, transcriptomic, and metabolomic approaches (11), the complexity of animal-associated microbiomes and the inability to culture and genetically-manipulate many symbionts make it difficult to study the precise molecular mechanisms that establish specific relationships.

The binary symbiosis between genetically-tractable *Vibrio fischeri* and the Hawaiian bobtail squid *Euprymna scolopes* serves as a model system to study the molecular interactions underlying microbiome assembly (11–16). The squid hatch aposymbiotically (without symbiont) and are colonized by *V. fischeri* in a multi-step process that leads to the specific recruitment of the symbiont from a marine environment in which the bacteria are < 0.1% of the bacterioplankton (14, 17). The symbionts are housed in the dedicated light organ (LO) within the squid’s mantle cavity, where they generate light that the host uses for counterillumination to hide its shadow while hunting at night (18). The host provides the symbionts with a protected niche, nutrients, and oxygen (15). Once the symbiosis is irreversibly established in juvenile squid, a daily cycle proceeds where 90-95% of the bacteria are expelled from the LO at dawn. The remaining symbionts grow during the day until they fill the LO, and at night the dense population of symbionts provides light (19). Because the aposymbiotic hatchlings can be cultured in the lab and infected with genetically-tractable *V. fischeri*, colonization experiments can be performed to study the molecular factors that play a role during this process (12, 16, 20). In addition, the translucent nature of the LO in squid hatchlings allows for visualization of the colonization process by microscopy (21–23).

Microbe-host signaling mechanisms and developmental transitions ensure specificity during colonization (14, 17). Upon detection of bacterial-derived peptidoglycan, the ciliated appendages on the surface of the LO secrete mucus that traps bacteria circulating within the mantle cavity (13, 24, 25). In the mucus field, *V. fischeri* bacteria bind to cilia and form aggregates by expressing symbiosis polysaccharide (*syp*) genes, a locus of 18 structural and regulatory genes whose products contribute to biofilm formation (26, 27). Approximately 3-4 hours post-inoculation, bacterial aggregates migrate through the host mucus toward the pores that lead into the LO ducts (25). While the initial migration is independent of flagellar motility (28), at the pores squid-produced chitin oligosaccharides serve as a chemoattractant to direct the symbiotic bacteria into the host crypts (21). Motility and chemotaxis are required for colonization, and strains with mutations in genes required for these processes—such as *cheA*, *flrA*, and *rpoN*—are unable to successfully colonize the squid LO (21, 28). Once within the LO *V. fischeri* generates light through expression of the *lux* operon in a quorum sensing-dependent process (29). Symbionts that fail to produce luminescence, such as strains with mutations in the autoinducer synthases *ainS* or *luxI*, or deletions of the *lux* operon, are unable to persist in the symbiosis (30, 31).

To identify novel colonization factors in *V. fischeri* our lab previously used a global transposon insertion sequencing approach (INSeq) to identify bacterial mutants that were depleted after 48 hours in the squid host (32). This approach successfully identified previously-known colonization factors, such as *rscS*, *rpoN, ompU*, various motility factors, and the *syp* biofilm locus, and in addition revealed 344 putative novel colonization factors. Twenty candidates were tested in competitive colonization assays of wild-type (WT) vs mutant strains and the results showed that nine factors had colonization defects. Some of the validated factors encompass roles in protein quality control (DnaJ and DegS) and copper detoxification (CopA and CusC), inner membrane proteins predicted to play a role in secretion of autotransporters (TamB/YtfN), and other poorly characterized factors (YdhC, YafD, and YhcB). This global approach was crucial in identifying putative colonization factors. However, further study is required to address which genes are true colonization factors, when they act during colonization, and how their products modulate the interaction with the host.

Approximately 32% of putative colonization factors identified by INSeq did not fall into a curated Clusters of Orthologous Groups (COGs) category, suggesting that the ability to interrogate the function of these colonization factors will reveal novel biology. Traditional genetic engineering techniques in *V. fischeri* are either random (transposon mutagenesis) or labor-intensive (plasmid-based allelic exchange) (32–34). We therefore considered approaches by which we could isolate mutants and examine phenotypes in a multiplexed fashion. One possible approach was to retrieve transposon insertions of interest from an arrayed library (35–37). A second approach we considered was to adapt a newly-developed method for transformation-mediated mutagenesis using linear DNA (38) with an in-frame barcoding strategy to facilitate precise mutations. This latter option was attractive in that we hoped that it would limit the effects of polar mutations and provide a set of defined deletions that can be characterized by amplicon PCR. Barcode sequencing (BarSeq), in which each strain is uniquely labeled and identified using high-throughput next-generation sequencing, has been used successfully to track population dynamics in multiple systems, including in yeast genomic libraries, during *Vibrio cholerae* infection, and to track and phenotype laboratory-evolved *Escherichia coli* (39–43). Here, we describe an approach to generate barcode-tagged gene deletions in *V. fischeri* and perform high-throughput colonization experiments using BarSeq. We also describe the barseq python computational package used to analyze the results.

## Results

### Generation of barcoded gene deletions

To generate barcoded deletions of specific *V. fischeri* genes, we designed an approach that takes advantage of splicing-by-overlap extension PCR and *tfoX*-based natural transformation (Fig. 1) (38, 44–46). The first step uses PCR to amplify DNA fragments upstream and downstream of the gene targeted for deletion, fused to the left and right linker sequences, respectively (Fig. 1A). A separate PCR is performed with pHB1 as a template to generate the central fragment of DNA containing the linker sequences, a selectable marker (*erm*, conferring erythromycin resistance) surrounded by FRT sites, and the semi-random barcode sequence. The barcode is provided by the reverse primer, which contains a region of semi-randomized-sequence. The three resulting DNA fragments—upstream, central, and downstream—are then fused into one fragment via their overlapping linker sequences by SOE-PCR (46) and transformed into *V. fischeri* upon *tfoX* induction (44). Finally, the *erm* cassette is removed via FLP recombinase (45).

**FIG 1.**
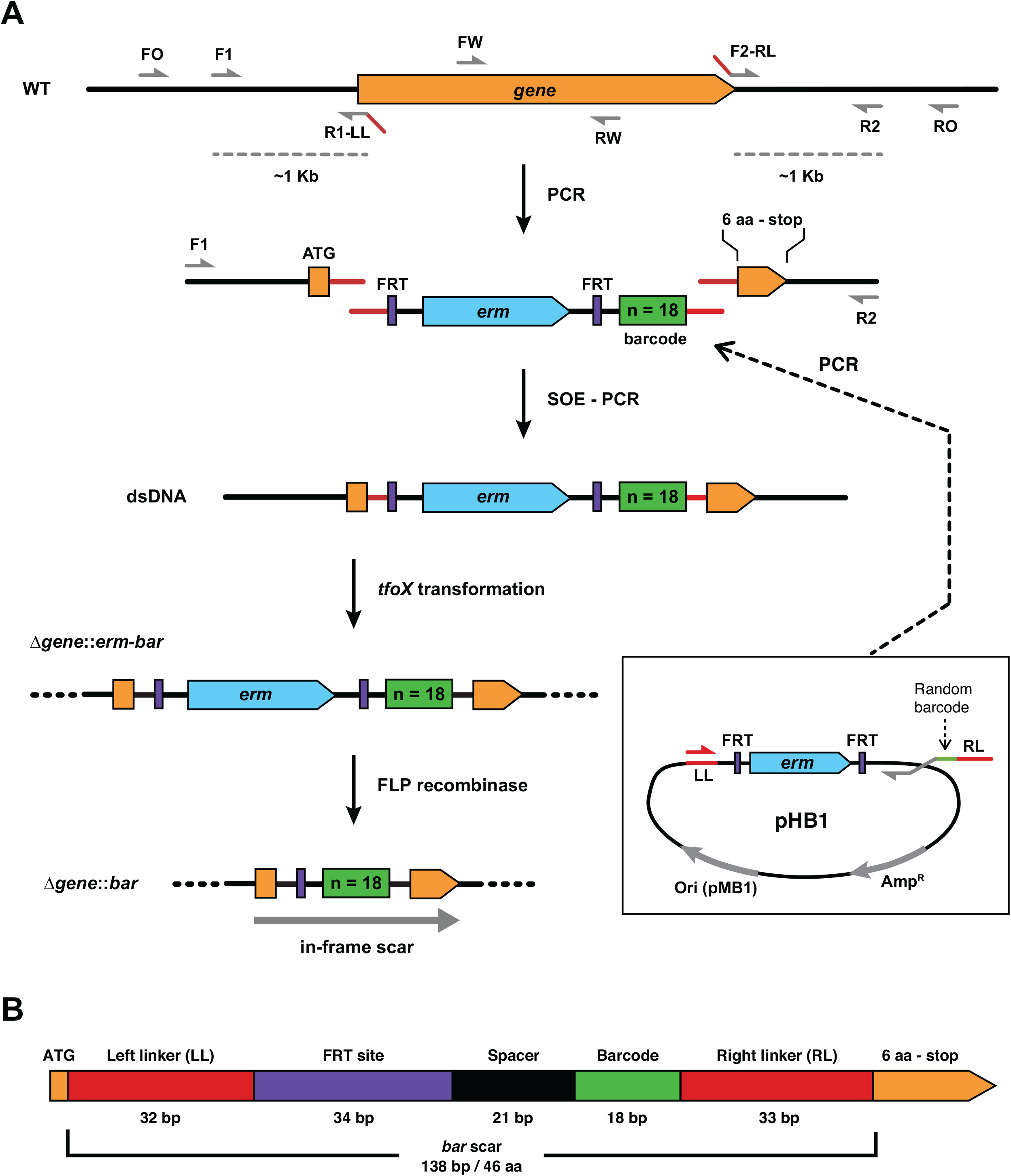
Approach for quickly generating barcode-tagged gene deletions in *V. fischeri*. (A) Schematic diagram (not to scale) of the process used to generate the barcoded deletions as described in the main text. Multiple primers are designed for use in PCR to generate the desired DNA molecules and screen/sequence for the correct deletion mutants as described in the Materials and Methods. (B) To scale schematic of the resulting *bar* scar containing the start codon, the left and right linker sequences (LL and RL), the FRT site that remains after removal of the *erm* cassette, a spacer sequence, the unique barcode, and the last 7 codons of the targeted open-reading frame (ORF). The barcode sequence is designed to lack in-frame stop codons, which results in an in-frame ORF together with the start codon and the last 7 codons of the targeted gene.

The resulting 138-bp deletion scar (*barcode* scar, or “*bar*” scar, Fig 1B) lies between the deleted gene’s first codon and last seven codons (i.e., the final six amino acid-encoding codons plus the stop codon). The scar is designed to be in-frame to prevent polar effects on gene expression when targeting genes within operons. The terminal codons were retained in case they contain a ribosomal binding site for downstream gene(s) (47). In addition to the barcode, the additional sequence in the scar includes left and right linker sequences that are shared among all of the mutants, which allows us to identify and quantify the abundance of each barcoded-strain using amplicon sequencing, while minimizing amplification bias by using common primers that amplify the same size product.

To test this new approach, we investigated the *copA* gene. Among Gammaproteobacteria, CopA is the main exporter of cytoplasmic copper and is the most widely conserved copper detoxification factor (48, 49). Although our laboratory previously demonstrated that *copA* is a squid colonization factor, its role in copper resistance has not been examined (32). We therefore targeted *copA* for deletion using our mutagenesis approach as a proof-of-concept and subsequently tested its role in copper resistance in *V. fischeri*. To ensure that the deletion process worked as intended, we used four sets of diagnostic PCR primers that would report on correct *erm* insertion, subsequent removal of the *erm* cassette by FLP recombinase, and absence of the targeted gene from the bacterial chromosome. PCR with various pairs of oligonucleotides that target the *copA* gene and its deletion constructs produced amplicons of the expected size in each strain (Fig. 2A). These results show that the *erm* cassette was successfully inserted into *copA* generating Δ*copA*::*erm-bar*, and subsequently removed by FLP recombination to generate the in-frame deletion scar in Δ*copA*::*bar*. Furthermore, sequencing of the deletion scar for several Δ*copA*::*erm-bar* candidates showed that after a single round of mutagenesis, multiple uniquely barcoded deletion strains were generated (Fig. 2B). These results demonstrate that our deletion method is successful in generating uniquely barcoded mutant strains of *V. fischeri*.

**FIG 2.**
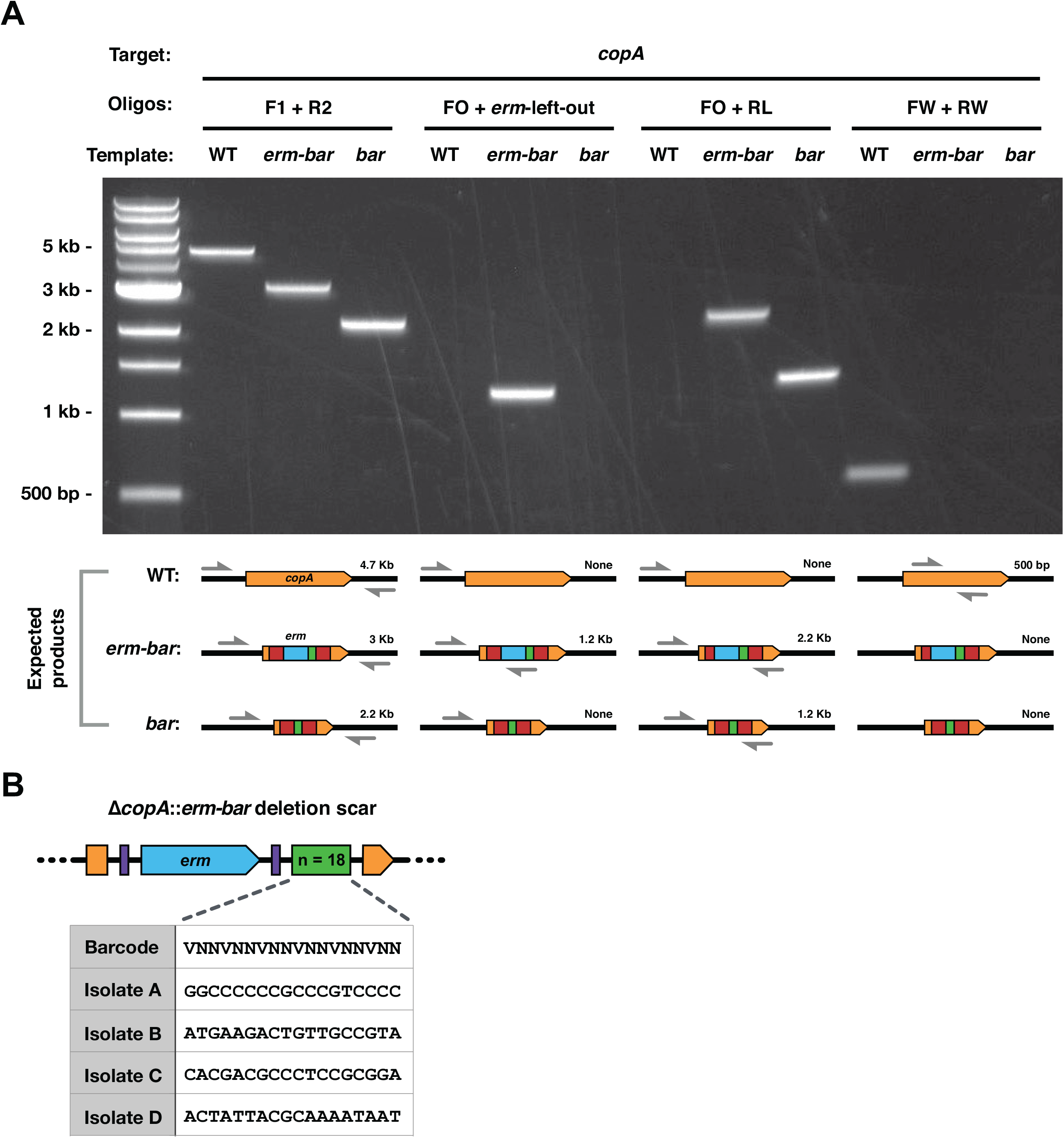
Evaluating the genotype of a *copA* deletion strain. (A) Representative 1% agarose gel showing the products generated by PCR when using the specified primer pairs and templates. DNA ladder is the 1 kb Plus DNA Ladder (New England BioLabs). (B) Table showing several unique barcode sequences within the Δ*copA*::*erm-bar* deletion scar of various deletion candidates that were generated from a single round of mutagenesis. Diagram is not to scale.

### The presence of a barcode within a gene deletion does not alter mutant phenotypes

To test that the barcoded scar does not affect the mutant phenotypes, we measured the copper sensitivity of strains deleted for *copA* using various methods. In addition to the mutants generated using our deletion approach (Δ*copA*::*erm-bar* and Δ*copA*::*bar*) we constructed a deletion of *copA* using plasmid-based allelic exchange (Δ*copA*) (33) and obtained a *copA* transposon mutant isolated from our previous study (Δ*copA*::Tn*erm*) (32). We then tested the growth of these various *copA* mutants in the presence of varying amounts of copper. Our results show that, regardless of the mutagenesis method, the growth of *copA* mutants is similarly impeded in the presence of copper, with the severity of the growth defect increasing in proportion to the concentration of copper: at 0.2 μM Cu^2+^ the *copA* mutants were able to grow slightly, whereas at 20 μM Cu^2+^ these strains were unable to grow (Fig. 3A). This is in contrast to the WT strain that achieved the same growth yield regardless of the concentration of copper present. The Δ*copA*::*erm-bar* and Δ*copA*::*bar* mutants showed the same degree of copper sensitivity (Fig. 3A).

**FIG 3.**
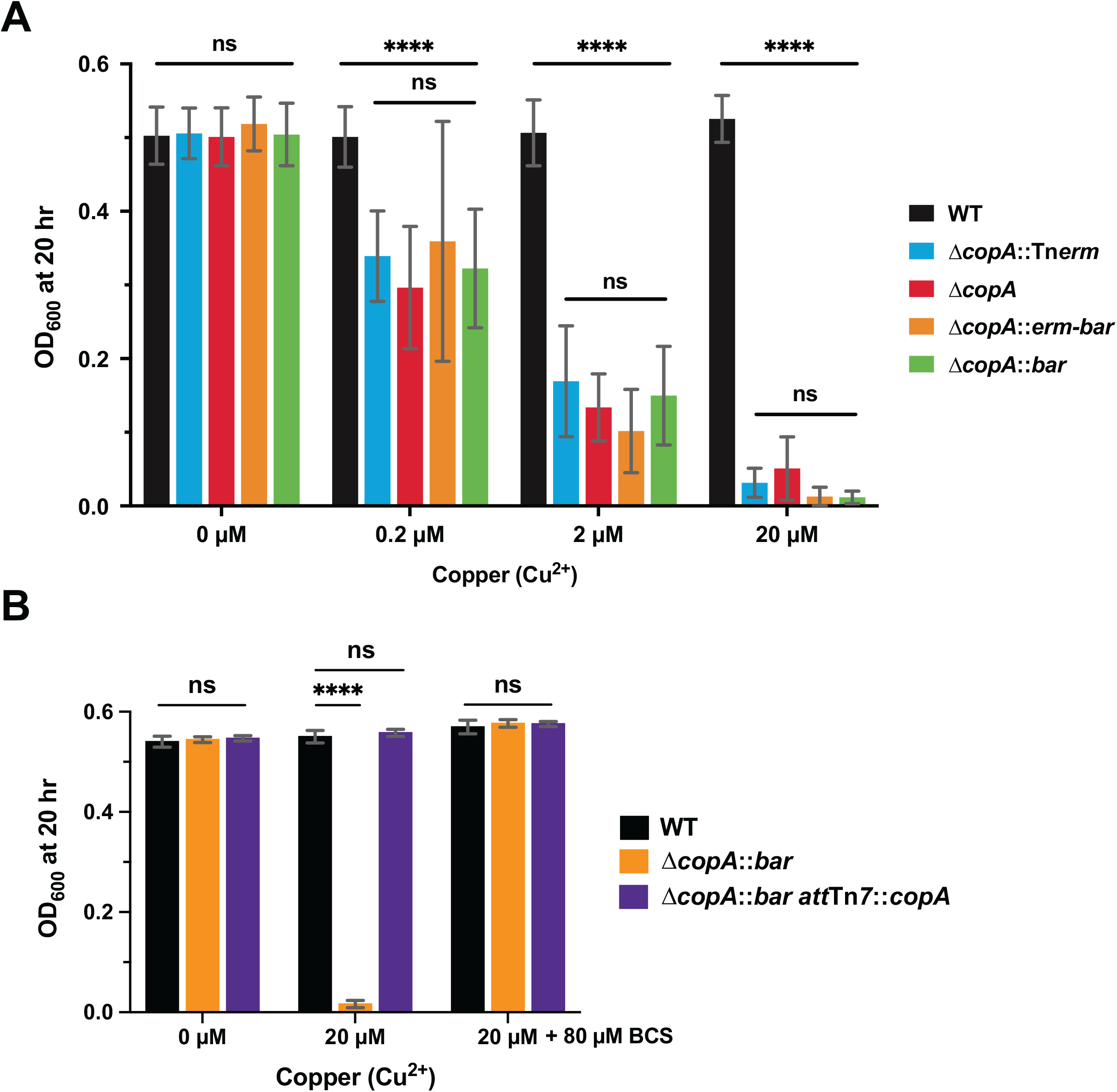
Assaying the phenotype of strains deleted for copper-resistance factors. (A and B) Bar graphs showing the average OD_600_ of the indicated Δ*copA* mutants after 20 hr of growth in the presence of the indicated amounts of copper and/or bathocuproinedisulfonic acid (BCS). (A) Error bars represent the standard deviation of the mean (*n* = 3). (B) Data are from two independent replicates (*n* = 2). Statistical analysis was performed using a Two-way ANOVA test. **** *P* < 0.0001.

To corroborate that the observed growth defects were due specifically to excess copper, we measured the growth of Δ*copA*::*bar* in the presence of copper, with and without the copper chelator bathocuproinedisulfonic acid (BCS). As expected, Δ*copA*::*bar* was unable to grow in the presence of 20 μM Cu^2+^, whereas the WT is unaffected (Fig. 3B). However, growth of Δ*copA*::*bar* in the presence of copper was rescued by addition of 80 μM of BCS (Fig. 3B), suggesting that free copper is indeed responsible for the observed lack of growth in the mutant. To verify that the absence of CopA was responsible for susceptibility to copper toxicity, we complemented *copA* at the chromosomal *att*Tn*7* site in the Δ*copA*::*bar* strain and observed that growth was rescued in the presence of copper (Fig. 3B). Based on these results, we conclude that CopA is required for resistance to copper in *V. fischeri*, consistent with its function in other Gammaproteobacteria (48).

To further test our deletion method, we generated mutants in multiple genes required for *V. fischeri* motility—*rpoN*, *flrA*, and *flaA*—and tested the resulting strains’ motility phenotypes on soft agar plates (28, 50, 51). We also included a WT strain tagged with the deletion scars at the *att*Tn*7* site (WT-1) and *copA* mutants as controls. While motility of WT *V. fischeri* resulted in a migration disc with a diameter of 26 mm from the inoculation point on soft agar plates, deleting *flaA* resulted in a drastic reduction in migration (9 mm), while deleting either *flrA* or *rpoN* resulted in no motility (1.5 mm) (Fig. 4). We observed that both the *erm-bar* and *bar* versions of the gene deletions displayed equivalent phenotypes, showing that the strains behave as null alleles regardless of whether the scar contains the *erm* cassette (Fig. 4). Both WT-1 and *copA* strains have motility comparable to WT, showing that motility defects are due to the deleted loci and not to the insertion of the deletion scars.

**FIG 4.**
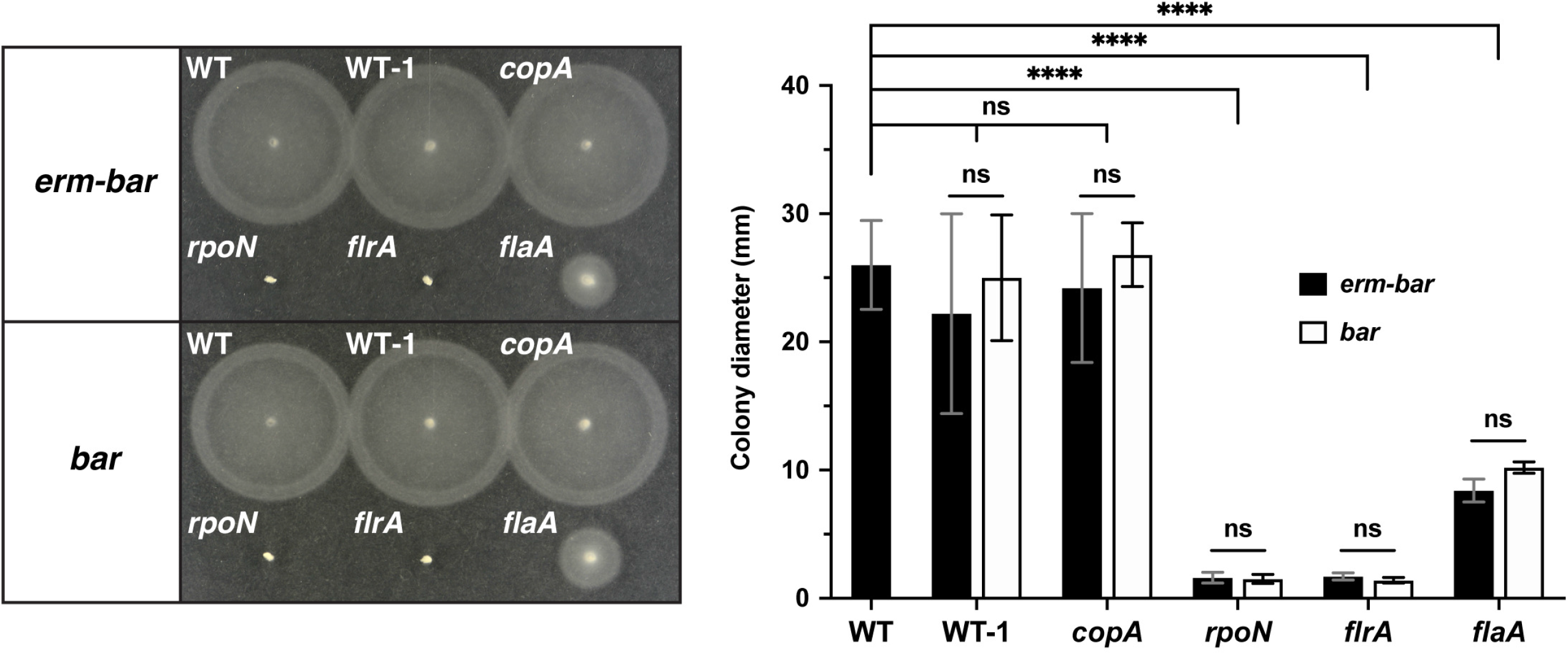
Assaying the phenotype of strains deleted for motility factors. Representative TBS agar trays showing the migration of strains from the inoculation point after incubation at 28°C for 4 hr. WT is MJM1100 (ES114), while WT-1 represents the *att*Tn*7*-marked MJM1100 strain with barcode 1 (either WT::*erm*-*bar*1 or WT::*bar*1). Bar graph shows the quantified data from five independent replicates with error bars showing the standard deviation of the mean (*n* = 5). Statistical analysis was performed using a Two-way ANOVA test. **** *P* < 0.0001.

### Removing the erythromycin-resistance cassette minimizes polar effects of the barcoded deletions

While the presence or absence of the *erm* cassette does not prevent deletion strains from manifesting the corresponding phenotypes (Fig. 3A and 4), we were concerned about polar effects on downstream gene expression upon insertion of the 1,049 bp heterologous *erm* cassette (52–54). To test the effect of the *erm* cassette on downstream gene expression we used reverse transcriptase quantitative PCR (RT-qPCR) to measure expression of genes immediately upstream and downstream of a targeted gene deletion for three different predicted operons. In each case, we measured the ratio in expression levels of the downstream vs upstream genes in the mutant, normalized to the ratio in WT *Vibrio fischeri* (defining this normalized value as the “polarity ratio”). For both *rpoN* and *cheA*, deletion scars of either *erm-bar* or *bar* resulted in negligible changes in the polarity ratio (Fig. 5). In contrast, the polarity ratio of Δ*cusA*::*erm-bar* was 26-fold higher than WT, whereas removal of the *erm* cassette to form the in-frame deletion scar restored the polarity ratio to basal levels (Fig. 5). We conclude that, in at least some cases, *gene*::*bar* deletion scars can alleviate collateral effects on flanking genes that are caused by inserting an antibiotic-resistance cassette.

**FIG 5.**
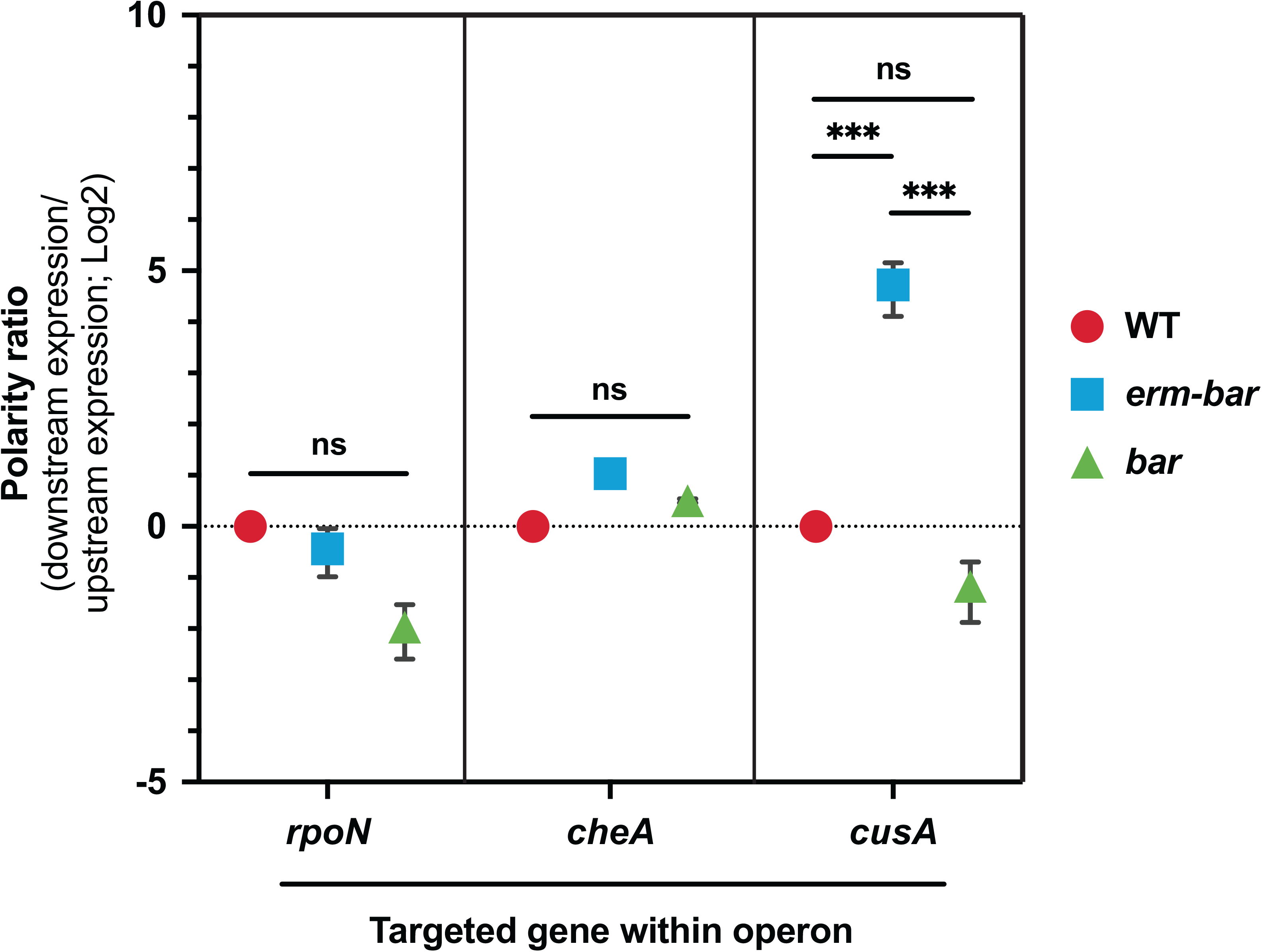
The *gene*::*bar* deletion scar reduces polar effects on gene expression introduced by the *erm* cassette. Graph showing the polarity ratio (expression of the downstream gene / expression of the upstream gene; relative to the indicated gene deletion target) for the indicated gene constructs within their respective predicted operons as measured by RT-qPCR. Statistical analysis was performed using a Two-way ANOVA test. *** *P* < 0.001.

### Development of a computational pipeline to analyze *V. fischeri* BarSeq data

With the ability to quickly generate precise barcoded deletions, we next sought to compete the Δ*gene*::*bar* deletions en masse during host colonization. We therefore developed a BarSeq sample preparation protocol, and an accompanying computational package to analyze the data (Fig. 6). To accomplish this, we mixed barcoded strains to generate an input library (i.e., a synthetic microbiome). This library was then used to inoculate media and/or squid hatchlings, which were then sampled at the desired time points. Samples were then processed to extract gDNA, and PCR was performed with dual-index Illumina sequencing primers to obtain dsDNA fragments containing the barcoded deletion scars. The resulting library was then sequenced on an Illumina MiSeq and demultiplexed based on the unique dual indexes (55). The resulting sequence data was then analyzed using the barseq package, which identifies and counts the barcodes present in the samples, assigns strain identity, normalizes strain counts, and calculates relative frequency and the competitive index (CI) for each strain within the samples. The BarSeq protocol provides a streamlined and effective way to measure population dynamics throughout squid colonization.

**FIG 6.**
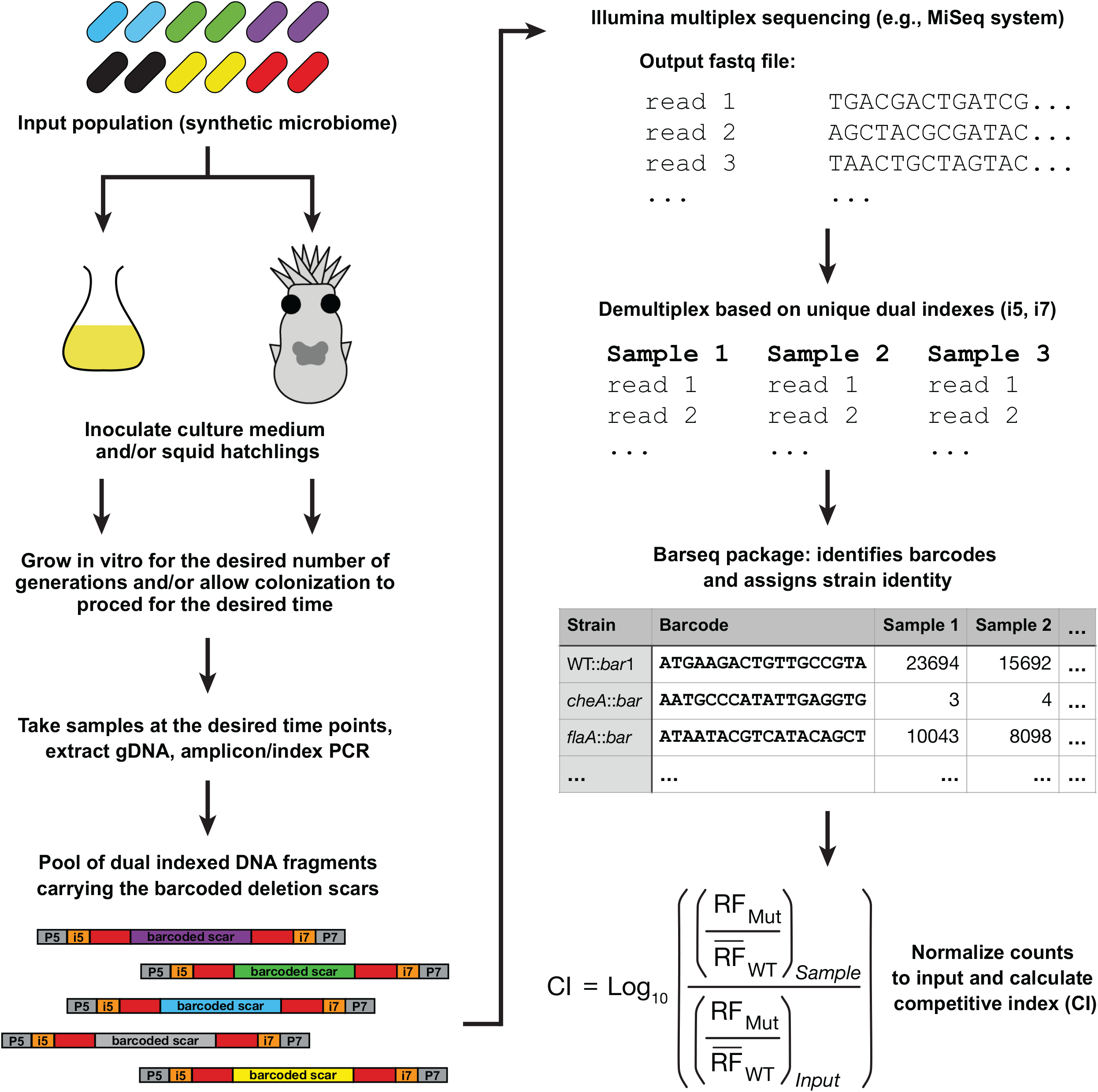
Overview of BarSeq experiments and computational package. Methodology and software for performing BarSeq experiments as described in the main text. An input population is used to inoculate squid or media, and samples are taken at the times of interest for gDNA extraction and processing to be sequenced by Illumina sequencing. The i5 and i7 segments are the index sequences in the dual indexed DNA fragments, whereas the P5 and P7 sequences are sequencing adapters for the MiSeq flow cell (details in Table S2). Sequencing reads are analyzed by the barseq package to obtain counts for individual strains in a sample based on their unique barcodes. Those counts are then used to calculate the relative frequencies of individual strains at each timepoint and the competitive index (CI) as described in Materials and Methods.

### BarSeq enables sensitive multiplex competition experiments

To test our BarSeq protocol in tracking individual strains within a population, we performed an *in vitro* competition and a competitive colonization experiment using an input library of seven different barcoded strains mixed in an equivalent ratio. In addition to several mutant strains, we included three WT::*bar* strains that had the *bar* scar inserted at the Tn*7* site that could be similarly tracked using amplicon sequencing but without affecting the phenotypes of the strains (WT-1, WT-2, and WT-3 in Fig. 7). After 15 generations of growth *in vitro*, the proportion of most strains relative to the WT::*bar* strains remained stable except for *flrA* and *rpoN*, which were 4-fold higher and lower, respectively, when compared to WT::*bar* (Fig. 7A). In contrast, following 48 h of squid colonization—which corresponds to approximately 15 bacterial generations (32)—we observed reduced levels of the *flaA* flagellin mutant and severely reduced levels of the *flrA*, *rpoN*, and *cheA* strains, all of which were near the limit of detection (Fig. 7B). This result is consistent with their previously known roles as necessary factors for squid colonization, although we did observe higher levels of *flaA* in the competitive colonization than are observed when a transposon insertion is competed against wild-type (28, 50, 51, 56). We note that there was relatively little variability among the WT::*bar* strains in the analysis, whereas the 4-5 log scale in which to identify colonization defects provided a substantially greater range over which to identify and refine colonization phenotypes *in vivo* (Fig. 7B). Taken together, these results show that our method for targeted barcoded deletions, multiplex squid colonization, and analysis by BarSeq allows for reproducible competition experiments *in vitro* and *in vivo* with high sensitivity.

**FIG 7.**
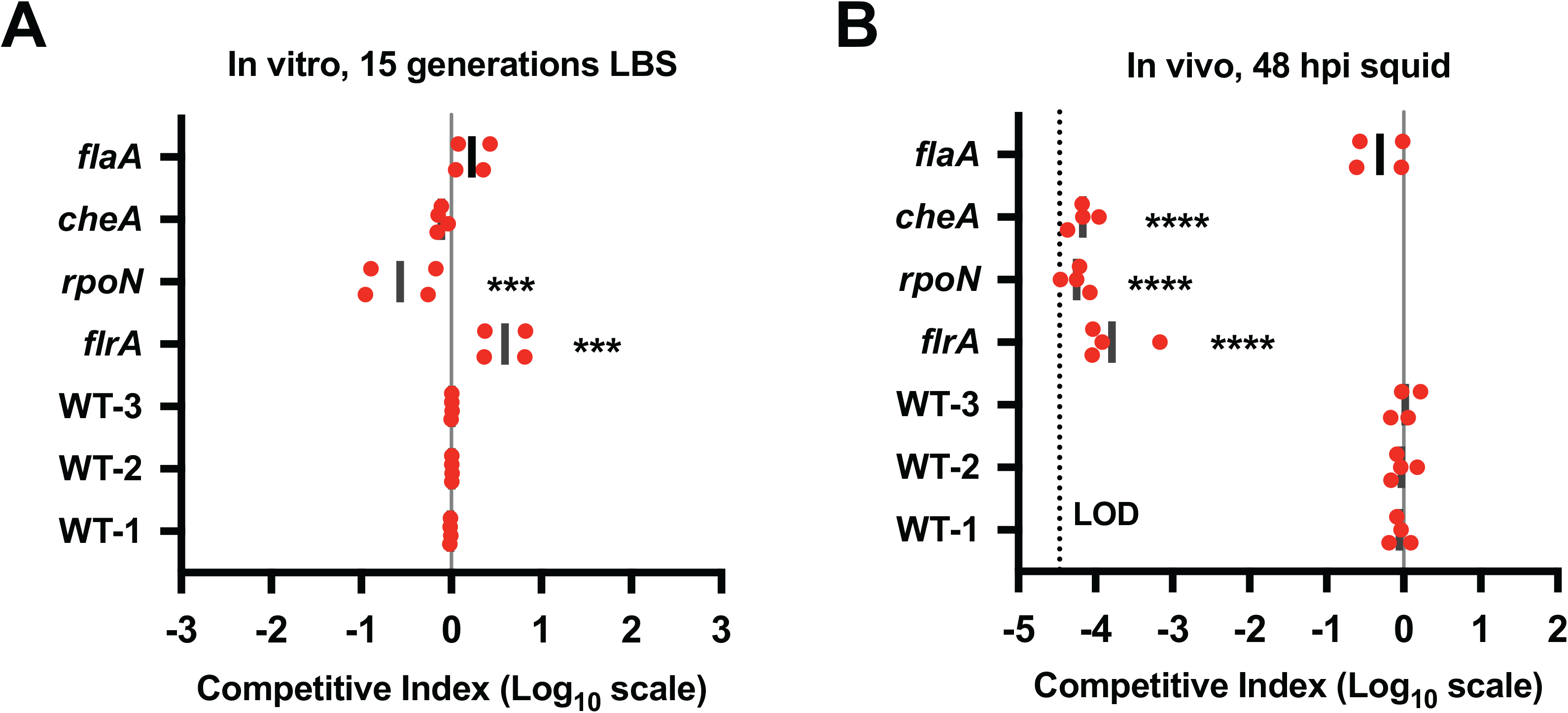
BarSeq enables high-throughput competition experiments. (A and B) Graphs show the mean competitive index (CI) on a Log_10_-scale for each barcoded strain in the population using the WT strains as controls as described in Brooks et al. (79) after (A) 15 generations *in vitro* in LBS and (B) 48 hr post-squid inoculation (hpi). WT is MJM1100 (ES114). WT-1 represents the *att*Tn*7*-marked MJM1100 strain with barcode 1 (WT::*bar*1), and similarly for WT-2 and WT-3 for barcodes 2 and 3, respectively. LOD = limit of detection for the experiment (3.39 × 10^−5^). Each symbol represents one biological replicate. Statistical analysis was performed using a One-way ANOVA test comparing each strain to WT-1. *** *P* < 0.001, **** *P* < 0.0001.

## Discussion

In this study, we developed a method to quickly generate gene deletions where the resulting strains are tagged with unique DNA barcodes. We demonstrated the utility of these strains in performing BarSeq high-throughput competitive colonization experiments and introduced a software package to analyze the resulting sequencing data. BarSeq provides a sensitive method to track population dynamics of squid colonization by *V. fischeri*.

### Generation of targeted, barcoded gene deletions that minimize effects on neighboring genes

Our approach builds upon previous SOE-PCR/*tfoX* mutagenesis techniques to incorporate a unique barcode in each deletion strain, which enables high-throughput experiments via barcode sequencing (BarSeq). Since BarSeq relies on amplicon sequencing, library preparation is straightforward and allows for a large number of samples to be processed in parallel. The method we have employed to design the barcode and flanking sequences was planned to minimize disruption on flanking genes. Expression of bacterial genes is frequently organized by their genetic arrangement into operons where expression of operon members is driven by a common promoter (57). However, given that some regulatory regions overlap in neighboring genes, deletion of one gene can alter the expression level of another nearby cistron. These off-target effects on gene expression could obfuscate the analysis of experimental results. Similar to the approach used by Baba et al. (53), our deletion approach reduces off-target effects on gene expression by ensuring the formation of an in-frame open reading frame within the deletion scar and including several codons at the end of the deletion target where the ribosome binding site of downstream genes is frequently located (Fig. 1B and Fig. 5).

### BarSeq enables detailed studies of the molecular mechanisms that result in establishment of the *Vibrio*-squid symbiosis

Using an INSeq screen, our lab previously identified 344 putative novel squid colonization factors in *V. fischeri* (32). Our deletion approach, combined with BarSeq, will enable the high-throughput characterization of these factors during squid colonization by allowing multiplexing of colonization factor mutants and tracking of individual strains. By enabling a more precise study of colonization factors, BarSeq has several potential applications.

BarSeq can be applied to the study of strain variation and evolution of colonization mechanisms in the *Vibrio*-squid symbiosis. *V. fischeri* strain variation is an important consideration when studying the mechanisms of colonization of the squid LO (58, 59). Previous studies have shown that multiple strains can co-colonize the squid LO and that they do so at different rates (60–62). More recent studies have focused on deciphering the specific mechanisms that result in differing colonization behavior (63, 64). The barcode-tagged mutagenesis method presented here can be applied to generate uniquely-tagged WT or mutant strains of the various phylogenetically-distinct *V. fischeri* strains and assayed in multiplexed format during squid colonization using BarSeq. We have already successfully used our SOE-PCR/*tfoX* mutagenesis approach to make targeted deletions in the ancestral strain SR5, showing that this method is applicable to *V. fischeri* strains that are evolutionarily distant to the frequently used ES114 strain (63).

BarSeq can also be used in directed evolution experiments to examine the functional evolution of colonization factors. Directed evolution experiments have recently been applied to study colonization factors in *V. fischeri* (65). The ease of tracking large numbers of individual *V. fischeri* strains using BarSeq could enable tracking of strain lineages in long-term evolution experiments, as has been conducted in other organisms (39, 42, 43).

### Phenotypes of *rpoN* and *flrA* mutants during competitive growth in media

Both *rpoN* and *flrA* deletion strains showed a statistically significant 4-fold decrease and increase, respectively, during competitive growth in media compared to WT (Fig. 7A). Due to the nature of the factors they encode, the observed growth defects are likely due to changes in energetic and nutritional requirements when *rpoN* and *flrA* are deleted. The *rpoN* gene encodes the alternative σ^54^ factor that is responsible for expression of various systems involved in squid colonization, including Syp biofilm formation, flagellar motility, and luminescence (51, 66, 67). Therefore, it is not surprising that deleting the gene encoding σ^54^ has pleiotropic effects on gene expression due to misregulation of the RpoN regulon and could reduce the ability of the mutant strain to effectively compete for growth *in vitro*, though further experiments are necessary to define the precise mechanism for the defect. FlrA is the σ^54^-dependent transcription factor that activates expression of the flagellar biosynthesis cascade and is required for motility and squid colonization (50). The high energetic cost of expressing all genes related to flagellar biosynthesis (68), which in *V. cholerae* requires FlrA-dependent regulation of 52 genes (69) and in *V. fischeri* between 39 and 131 genes (28), is consistent with the observed increase in growth of the *flrA* deletion strain compared to WT during competitive growth in media (Fig. 7A). Nonetheless, even though the changes observed in competitive growth of the *rpoN* and *flrA* mutants *in vitro* are in opposing directions (less vs more growth, respectively), both are severely defective in squid colonization (Fig. 7B). Future experiments using BarSeq to probe bacterial growth *in vitro* and during colonization have the potential to elucidate heretofore hidden phenotypes.

### Discrepancy in *flaA* colonization efficiency measured by BarSeq vs traditional competitive squid colonization experiments

In our BarSeq experiment, the known colonization factor *flaA* only shows a small (~2-fold) colonization defect after 48 hr post-squid inoculation (Fig. 7B). In contrast, Millikan and Ruby showed that a *flaA* deletion made by insertion of a Kan^R^-cassette is severely defective during competitive colonization against WT *V. fischeri* (56). Using confocal microscopy, their work showed that LO colonization by *flaA* is delayed compared to WT by ~8 hr. However, because our competitive colonization experiment using BarSeq was done at 48 hours post-inoculation, this delayed colonization is not enough to explain the observed discrepancy. Previous work has shown that the concentration of *V fischeri* in the inoculum can affect the number of different strains that can co-colonize the squid LO (70). This raises the possibility that the inoculum amount or the ratio of strains within the synthetic microbiome might affect the observed colonization defect. To address this, future experiments should examine how inoculum amount and the ratio of mutant strains to WT within a synthetic barcode-tagged population affects colonization efficiency for the different strains in the population.

In summary, we provide a new method for constructing barcoded deletions of *V. fischeri* genes; demonstrate the utility of this method for generating in-frame deletions and discovering new functions of squid colonization factors; and combine this approach with a computational tool to conduct multiplex animal colonization assays using barcode sequencing (BarSeq).

## Materials and Methods

### Bacterial strains, growth conditions, plasmids, and primers

Bacterial strains used in this study are listed in Table 1, with Table S1 containing an Extended Table 1 showing the oligos used to generate the specified barcode-tagged gene deletions. Plasmids are listed in Table 2, and DNA oligonucleotides are listed in Table S2. DNA oligonucleotides were synthesized by Integrated DNA Technologies (Coralville, IA), and Sanger DNA sequencing was performed through the University of Wisconsin–Madison Biotechnology Center. *Escherichia coli* strains were grown in Luria-Bertani (LB) medium [per liter, 25 g Difco LB broth (BD), in distilled water] at 37°C with aeration. Unless otherwise indicated, *V. fischeri* strains were grown in Luria-Bertani salt (LBS) medium [per liter, 25 g Difco LB broth (BD), 10 g NaCl, and 50 ml 1 M Tris buffer, pH 7.0, in distilled water] at 25°C with aeration. When necessary, growth media was solidified by adding 15 g Bacto agar (BD) per liter. For growth of *V. fischeri*, antibiotics (Gold Biotechnology) were added at the following concentrations: 5 μg/ml erythromycin, 5 μg/ml or 2.5 μg/ml chloramphenicol as indicated, and 100 μg/ml kanamycin. For *E. coli* the antibiotic concentrations used were 100 μg/ml carbenicillin, 25 μg/ml chloramphenicol, and 50 μg/ml kanamycin. The *E. coli* strain π3813 containing pKV496 is a thymidine auxotroph and was grown in LB with 50 μg/ml kanamycin supplemented with 0.3 mM thymidine (38, 71).

**Table 1.**
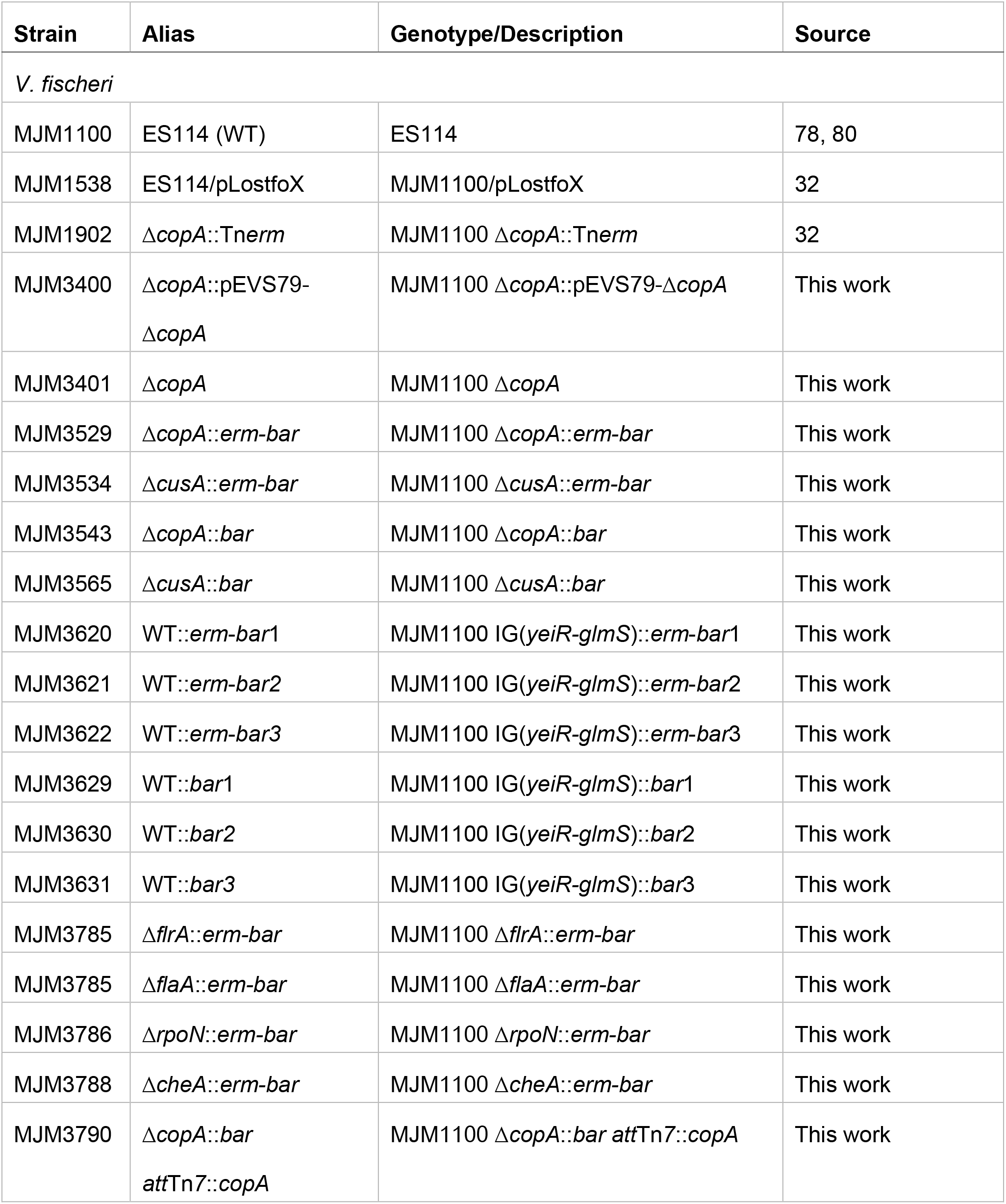
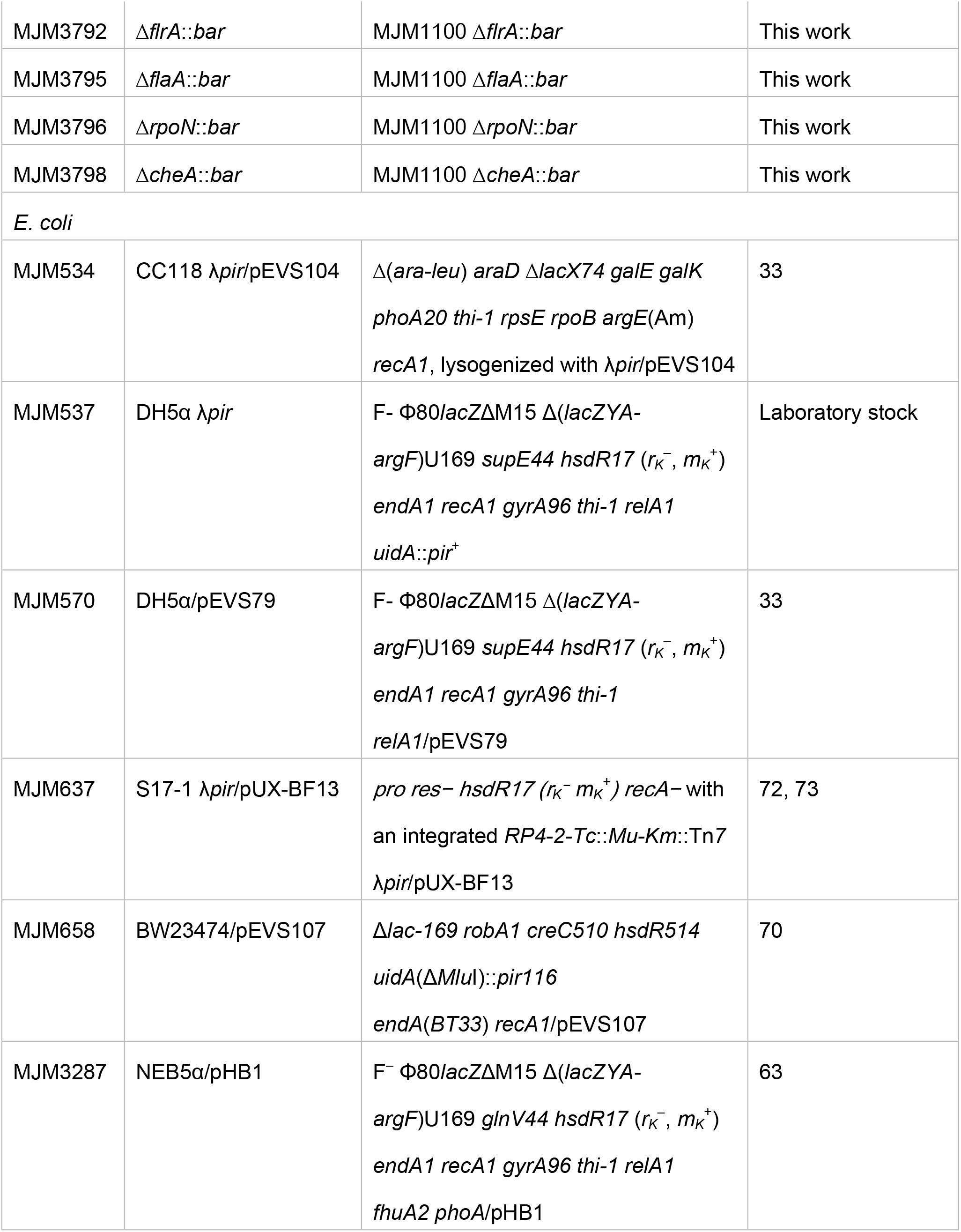
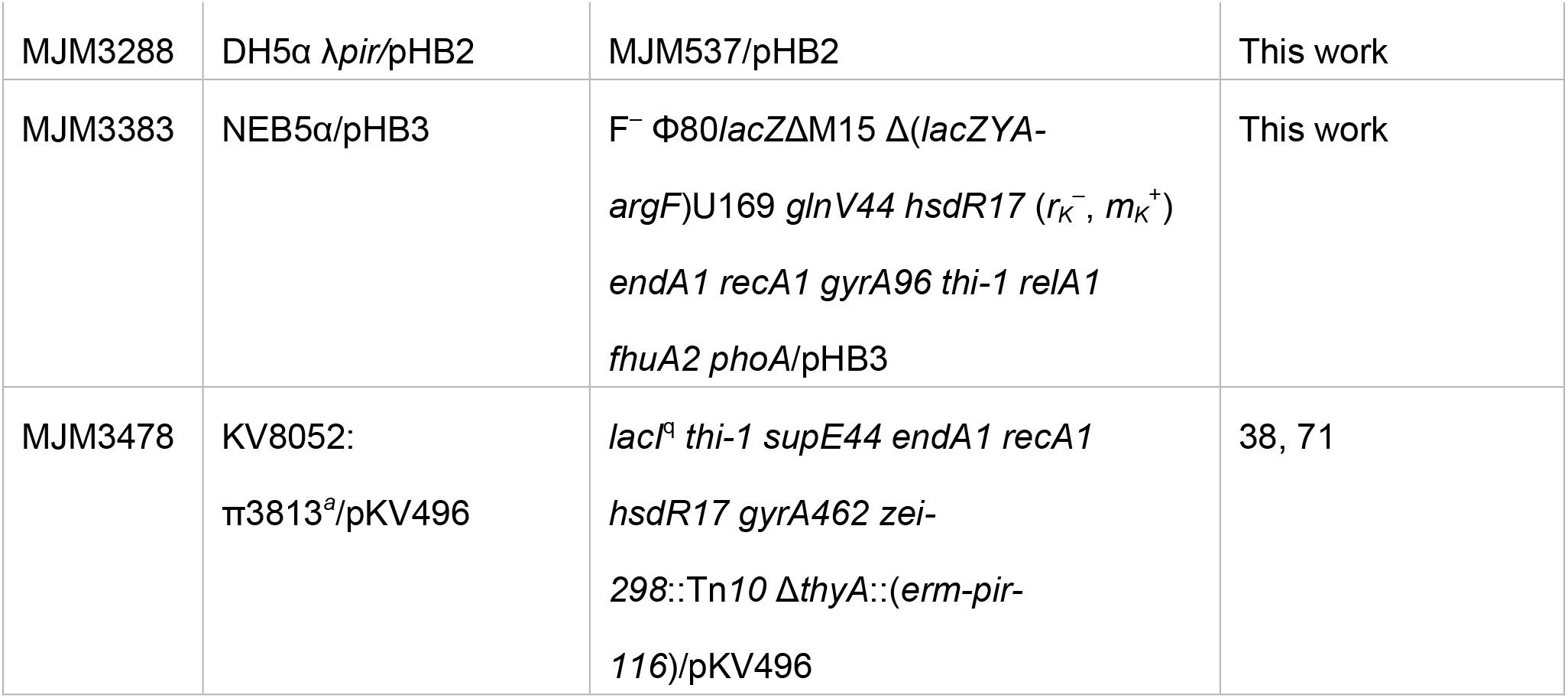
Bacterial strains. ^*a*^Thymidine auxotroph, growth conditions in Materials and Methods. N/A = Not applicable.

**Table 2.** Plasmids.

| Plasmid | Relevant properties | Source |
| --- | --- | --- |
| pEVS79 | Vector backbone for deletion construct via allelic-exchange, Cam<sup>R</sup> | 33 |
| pEVS104 | Conjugation helper plasmid, Kan<sup>R</sup> | 33 |
| pEVS107 | mini-Tn7 mobilizable vector, Erm<sup>R</sup> (transposon), Kan<sup>R</sup> | 70 |
| pKV496 | pEVS79 containing the FLP recombinase, Kan<sup>R</sup> | 38 |
| pLostfoX | *tfoX* overexpression vector, Cam<sup>R</sup> | 44 |
| pUC19 | Cloning vector, Carb<sup>R</sup> | Laboratory stock |
| pUX-BF13 | Tn7 transposase helper plasmid (*tns* genes), Carb<sup>R</sup> | 72 |
| pHB1 | pUC19 containing the LL-FRT-*erm*-FRT-spacer sequence in the HindIII/BamHI site | 63 |
| pHB2 | pEVS107 containing *copA* (including 191 bp upstream and 321 bp downstream of the *copA* ORF) at the Ascl site | This work |
| pHB3 | pEVS79 containing 1.6-kb upstream/1.6-kb downstream of *copA* | This work |

The unmarked deletion of *copA* in MJM1100 was made by allelic exchange as described previously (63). Briefly, 1.6 kb upstream (US) and 1.6 kb downstream (DS) sequences of *copA* were amplified by PCR using oligos HB44 and HB45, and HB46 and HB47, respectively, and were cloned using Gibson Assembly (NEBuilder HiFi DNA assembly cloning kit) into the linearized vector pEVS79 (linearized using oligos HB52 and HB53) (Table S2). The Gibson mix was transformed into NEB5α chemically-competent cells and selected on chloramphenicol. The resulting pEVS79-Δ*copA* candidates were screened using PCR with oligos HB54 and HB55 and confirmed by sequencing, generating pHB3, which was conjugated into *V. fischeri* MJM1100 (ES114) via triparental mating with MJM534, which contains the helper plasmid pEVS104 (33). Single recombinants of pHB3 into the chromosome were screened and selected by growth on chloramphenicol (MJM3400), and double recombinants by loss of the antibiotic resistance cassette and *copA* (MJM3401). The resulting constructs were verified by PCR and sequencing (Table S2).

The *copA* gene was inserted into the *att*Tn*7* site in the chromosome using pEVS107 (70). The *copA* gene including 191 bp US and 321 bp DS sequences was amplified by PCR using oligos HB27 and HB34, the product was digested with AscI, and cloned into the AscI site of pEVS107. The resulting plasmid, pHB2 (pEVS107-*copA*), was transformed into and maintained in *E. coli* DH5α λ*pir* cells and verified by PCR (Table S2) and sequencing. pHB2 was conjugated into Δ*copA* (MJM3401) via tetraparental mating with donor MJM3288 (DH5α λ*pir*/pHB2), helper strains MJM637 (S17-1 λ*pir*/pUX-BF13) (72, 73) and MJM534 (CC118 λ*pir*/pEVS104) (33), and the recipient MJM3543 (Δ*copA*::*bar*), resulting in MJM3790 (Δ*copA*::*bar att*Tn*7*::*copA*). Candidates were confirmed by PCR (Table S2) and sequencing.

### Construction of barcode-tagged gene deletions

The deletion protocol demonstrated in Fig 1A is based on splicing by overlap extension PCR (SOE-PCR) and *tfoX* transformation (38, 44–46) to directly delete and tag targeted genes with a randomized sequence (barcode). Our protocol was in development prior to publication of the previous method (38), so while it is conceptually similar, the sequences of the linkers and primers are distinct. First, several oligos were designed specific to the targeted genes to amplify 1 kb of US (F1 and R1-LL) and DS (F2-RL and R2) DNA tagged with the left linker (LL) and right linker (RL) sequences, respectively, to screen the deletion scar via PCR (FO and RO), and to assay for the absence of the targeted gene (FW and RW) (Fig. 1A, Table S1, and Table S2). FW and RW were designed to amplify a fragment of 500-1,000 bp, depending on the size of the gene. The F1 and R2 oligos were designed to anneal 1 Kb US and DS, respectively, of the targeted gene. The R1 oligo was designed to anneal starting at the start codon of the targeted ORF going upstream, then the reverse complement of the LL sequence (LL reverse complement: 5’-CTGGCGAAGCATATATAAGAAGCTCGTCTCGT-3’) was attached to the 5’-end of the R1 oligo, resulting in R1-LL. The F2 oligo was designed to anneal at the last 7 codons (6 aa and stop codon) on the 3’-end of the targeted ORF going downstream, then the RL sequence (RL: 5’-GACTTGACCTGGATGTCTCTACCCACAAGATCG-3’) was attached to the 5’-end of the F2 oligo, resulting in F2-RL. The FO and RO oligos (<u>f</u>orward <u>o</u>utside and <u>r</u>everse <u>o</u>utside, respectively) were designed to anneal 500 bp away from the annealing sites of F1 and R2, respectively, and were used to probe the targeted genomic region for insertion of the desired deletion scar.

The middle dsDNA fragment containing the *erm* cassette flanked by FRT sites and the randomized barcode was obtained by PCR with Phusion Hot Start Flex 2X master mix (NEB; M0536L) and pHB1 as template, which contains the LL-FRT-*erm*-FRT-spacer sequences and was built as described previously (63), and oligos HB42 and HB154. Oligo HB154 is a reverse primer and contains the RL sequence, 18 bp of randomized sequence composed of 6 trimers of ‘NNB’ to prevent formation of stop codons (results in ‘VNN’ codons in the forward direction), and the spacer sequence (Table S2). The resulting 1,049 bp product containing LL-FRT-*erm*-FRT-spacer-random barcode-RL was purified by gel extraction using a QIAquick Gel Extraction Kit (Qiagen; 28706). The flanking 1 kb US and DS fragments for each targeted gene were then fused to this middle DNA fragment via the homology between the LL and RL sequences and using SOE-PCR with the F1 and R2 oligos, resulting in the 3 kb mutagenic dsDNA. The reaction mixture contained 10 ng of each the middle, US, and DS DNA fragments, 200 nM of the corresponding F1 and R2 oligos (Table S2), 1X Phusion Hot Start Flex Master Mix (NEB; M0536L) and H_2_O up to a total volume of 25 μl. SOE-PCR conditions were 98°C for 30 sec, 98°C for 5 sec, 60°C for 20 sec, 72°C for 1 min (30 cycles), with a final extension step at 72°C for 5 min.

The 3 Kb mutagenic DNA fragments were purified using a QIAquick PCR Purification Kit (Qiagen; 28106) and transformed into *V. fischeri* ES114 via natural transformation with pLostfoX (MJM1538) (32, 44) where the flanking sequences guide the barcoded *erm* cassette to substitute the targeted gene. Mutant candidates were selected on LBS-Erm5 and screened by PCR with oligo pairs F1/R2, FO/HB8, and FW/RW (as shown in Fig. 2A). The insertion of the *erm-bar* scar was confirmed by Sanger sequencing with primers HB8, HB9, HB42, and HB146, and the unique barcode sequence was recorded for each strain.

The final *bar* scars were made by triparental mating of donor MJM3478 (π3813/pKV496) (38) and helper strain MJM534 (CC118 λ*pir*/pEVS104) into recipient *V. fischeri* strains containing the *erm-bar* scar and selection on LBS-Kan100. Plasmid pKV496 contains the FLP recombinase that removes the *erm* cassette and fuses the two surrounding FRT sites into one, resulting in the final *bar* scar as shown in Fig. 1B. The plasmid was eliminated by growing the candidates on LBS without selection twice and selecting colonies that were Erm^S^ and Kan^S^. The *gene*::*bar* candidates were screened by PCR using oligo pairs F1/R2, FO/HB146 (RL), and FW/RW, and the deletion scar verified by Sanger sequencing using oligos HB42 and HB146. The barcode sequences were verified to match the barcode within the parental strains containing the *gene*::*erm-bar* scar.

The barcoded WT *V. fischeri* strains (WT::*bar*) were constructed using the same procedure as outlined above for the gene deletions but targeting a site next to the *att*Tn*7* site in the intergenic region of *yeiR* and *glmS*. The 1 kb US and DS arms were amplified using PCR with ES114 gDNA and oligo pairs HB239/HB240 and HB241/HB242. After SOE-PCR to form the mutagenic DNA and *tfoX* transformation, the WT::*erm-bar* candidates were screened by PCR with oligo pairs HB243/HB244 and HB243/HB8. Sanger sequencing was used to confirm insertion of the *erm-bar* scar and record the unique barcode sequences. Triparental mating as described above was performed to remove the *erm* cassette using pKV496. The *bar* scar was confirmed by PCR with HB243/HB146 and Sanger sequencing.

### Growth assays in the presence of copper

Colonies from freshly streaked LBS plates of the indicated *V. fischeri* strains were inoculated into 3 ml LBS with the appropriate antibiotics and grown for 8 hr at 25°C with shaking. Three microliters of the LBS cultures were subcultured into 3 ml Tris minimal medium [per liter, 500 ml DSW (2X), 50 ml 1 M Tris base, pH 7.5, 1 ml 5.8% K_2_HPO_4_, 1 ml 10 mM FeSO_4_, and 20 ml 10% *N*-acetylglucosamine (GlcNAc), in distilled water; DSW (2X) = 100 mM MgSO_4_, 20 mM CaCl_2_, 600 mM NaCl, and 20 mM KCl] and incubated at 25°C overnight for ≤ 16 hr. Overnight cultures were diluted to an OD_600_ of 0.5 in 200 μl, then 2 μl of 0.5 OD_600_ were transferred into 198 μl of fresh Tris minimal medium containing the appropriate amounts of copper and/or BCS in a 96-well plate. The plate was then incubated in a Synergy Neo2 Multi-Mode Microplate Reader (BioTek) at 25°C with OD_600_ measurements every 15 min for 20 hr. Copper stock solutions (100 mM) were prepared from copper (II) sulfate pentahydrate (CuSO_4_·5H_2_O; Sigma-Aldrich; 203165), and BCS stock solutions (50 mM) from bathocuproinedisulfonic acid disodium salt (Sigma-Aldrich; B1125).

### Motility assays

The indicated bacterial strains were streaked onto fresh LBS plates with the appropriate antibiotics and grown overnight at 25°C. Single colonies were picked with a sterile toothpick and deposited onto OmniTrays (Thermo Fisher Scientific; 242811) containing TBS Agar [per liter, 10 g Gibco Bacto Tryptone (Thermo Fisher Scientific; 211705), 50 ml 1 M Tris buffer, pH 7.0, 20 g NaCl, 8.63 g MgSO_4_, and 3 g Agar, in distilled water] by stabbing the toothpick into the media at a single spot. Trays were incubated at 28°C for 4 hr and the outer diameter of swimming cells was measured.

### Measuring polarity ratio via RT-qPCR

The indicated bacterial strains were grown in 3 ml LBS with the appropriate antibiotics and grown at 25°C overnight. On the day of the experiment, 15 μl of the overnight cultures were transferred into 3 ml of fresh LBS and growth was continued at 25°C with aeration. Samples were harvested at an OD_600_ of 0.2-0.4 (mid-log phase) by transferring 800 μl of culture into a 2 ml screw-cap tube containing 100 μl of a cold 95% EtOH– 5% Phenol solution that inactivates RNases (74). RNA extraction and RT-qPCR were performed as described previously (75). Briefly, cells were lysed in Tris-EDTA (TE) buffer (10 mM Tris-Cl, pH 8.0, 1 mM EDTA) containing lysozyme (Epicentre; R1804M) and 1% SDS. RNA was extracted using the hot phenol method (74) and digested with DNase I (NEB; M0303S).

cDNA was synthesized from 0.5 μg of total RNA using the iScript Advanced cDNA synthesis kit (Bio-Rad; 1725037) following the protocol 25°C for 5 min, 46°C for 20 min, and 95°C for 1 min. Quantitative PCR was performed using 1:10 dilutions of cDNA synthesis products with the iTaq Universal SYBR green supermix (Bio-Rad; 1725121) on a CFX Connect real-time PCR detection system (Bio-Rad). The qPCR protocol was 95°C for 30 sec, 95°C for 5 sec, 58°C for 30 sec (40 cycles), with a final melt curve analysis to ensure specificity in the reaction. The mRNA levels of *rpoD*, *lptB*, *hpf*, *cheZ*, *cheB*, *cusB* and *cusF* were measured using the oligo pairs listed in Table S2. Expression levels for each gene were normalized to *rpoD* and the mutants were normalized to WT using the 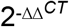 method (76). The polarity ratio of *rpoN*, *cheA*, and *cusA* was calculated as “expression of the downstream gene / expression of the upstream gene” using the respective flanking genes in each putative operon–*lptB*-*rpoN*-*hpf*, *cheZ*-*cheA*-*cheB*, and *cusB*-*cusA*-*cusF*. Operons were predicted using the BioCyc database for “*Aliivibrio fischeri*”, Strain ES114, version 24.1, which is based on the sequenced genome in Mandel et al. (77, 78).

### Barseq bioinformatic tool

To quantify barcodes within each sequenced sample, we developed barseq (https://github.com/mjmlab/barseq), a python package that identifies putative barcodes in the sequenced reads and matches them to a user provided barcode library. The program iterates through each sample and uses regular expressions to search within the reads for flanking sequences on the left (GCTCATGCACTTGATTCC; spacer sequence) and the right (GACTTGACCTGGATGTCT; right linker sequence) of the barcode (Fig. 1B), while also allowing for 18 random nucleotides that represent a candidate barcode. The putative barcode sequence is then mapped against the reference barcode library and increases the count for the matched strain. Barseq outputs a tab-delimited table with the barcode/strain counts for each of the samples analyzed.

### Barcode Sequencing and multiplexed competitive experiments

Cells of the indicated strains (Fig. 7) were grown in 3 ml LBS at 25°C overnight with aeration. The cultures were then diluted (1:80) into 3 ml fresh LBS and grown to mid-log phase (0.2 OD_600_). Equivalent ODs of cells from each strain [volume to mix calculated as Vol. (μl) = (1.25/OD_600_) X 50] were mixed, resulting in a multiplexed population with each strain present at a 1 to 1 ratio. A sample from this input library was harvested by collecting cells from 700 μl by centrifugation and storing the cell pellet at −80°C. The input library was then used to inoculate hatchling *Euprymna scolopes* squid at 5-9 × 10^3^ CFU/ml for 3 hr in FSIO (filter sterilized Instant Ocean) as previously described (20). Squid samples (n = 24, per replicate) were harvested at 48 hr post-inoculation and surface sterilized by storing at −80°C. Concurrently to squid colonization, the input library was competed for growth *in vitro* for 15 generations by diluting the library 1:181 into LBS, growing at 25°C with aeration back to the starting OD_600_, repeating this process once more, and harvesting samples as described above. Individual squid were homogenized in 700 μl of FSIO, 500 μl of each homogenate was mixed in a 50 ml conical tube, diluted 1:20 in 70% IO (Instant Ocean), and 50 μl plated onto LBS plates in triplicate. After a 17 hr overnight incubation at 25°C the bacterial colonies from each plate were scraped with a sterile cell scraper into 1 ml of 70% IO and collected by centrifugation. Cell pellets were stored at −20°C prior to DNA extraction.

Genomic DNA from the cell pellets was extracted and purified using the Qiagen DNeasy Blood and Tissue Kit (Qiagen; 69506) following the Gram-negative bacteria protocol, and was quantified using a NanoDrop spectrophotometer (Thermo Scientific). The barcoded scars were amplified together with dual-index Illumina sequencing primers (55). The reaction mixtures contained 50 ng of gDNA, 200 nM of each oligo (Table S2), 1X Phusion Hot Start Flex Master Mix (NEB; M0536L) and H_2_O up to a total volume of 50 μl. PCR conditions were 98°C for 30 sec, 98°C for 10 sec, 60°C for 10 sec, 72°C for 10 sec (20 cycles), with a final extension step at 72°C for 5 min. PCR products were visualized using a 2% agarose gel to confirm the dual-indexed amplicon of 231 bp and purified using a QIAquick PCR Purification Kit (Qiagen; 28106). Purified PCR products were quantified using a Qubit 3 fluorometer (Life Technologies) and pooled in equimolar amounts, and diluted to 4 nM. The pool was sequenced on an Illumina MiSeq using the 2 × 250 bp v2 kit with a 10% PhiX control following the manufacturer’s protocol (Illumina, Inc., San Diego, CA) and using custom primers developed from (55). The sequencing data was processed using the barseq python package to obtain strain counts per sample, and mutants that were in the input library but still being validated were removed from the dataset. The relative frequency (RF) for each strain in a sample was calculated, normalized to the RF in the input library and the average RF in the sample, and the competitive index (CI) was then calculated using the formula: CI = Log_10_[(RF_mutant_/Avg. RF_WT_)_Sample_/(RF_mutant_/Avg. RF_WT_)_Input_].

## Supporting information

Supplemental References

## Supplementary Tables S1-S3

- **Table S1. Expanded bacterial strains**. ^*a*^Thymidine auxotroph, growth conditions in Materials and Methods. N/A = Not applicable.
- **Table S2. DNA oligonucleotides.**
- **Table S3. Strain counts from barseq output.** Strain counts for individual strains within each sample was obtained using the barseq package. The samples for competitive squid colonization (48 hpi) were processed in triplicate as technical replicates. The counts in ‘_other’ represent sequence reads that contain the appropriate sequences flanking the barcode region but the barcode sequence does not match any present in the reference barcode library (as described in Materials and Methods).

## Acknowledgments

We thank Karen L. Visick for plasmid pKV496, and members of the Mandel lab for comments on the mutagenesis protocol and manuscript.

